# Mitochondrial OPA1 deficiency causes reversible defects in adult neurogenesis-associated spatial memory in mice

**DOI:** 10.1101/2021.11.16.468792

**Authors:** Trinovita Andraini, Lionel Moulédous, Petnoi Petsophonsakul, Cédrick Florian, Sébastien Lopez, Marlène Botella-Daloyau, Macarena Arrázola, Kamela Nikolla, Adam Philip, Alice Leydier, Manon Marque, Laetitia Arnauné-Pelloquin, Pascale Belenguer, Claire Rampon, Marie-Christine Miquel

**Affiliations:** Centre de Recherches sur la Cognition Animale (CRCA), Centre de Biologie Intégrative (CBI), Université de Toulouse, CNRS, UT3, France; Department of Physiology, Faculty of Medicine, Universitas Indonesia, Jakarta, Indonesia

**Keywords:** hippocampus, adult neurogenesis, DOA, pattern separation, physical exercise

## Abstract

Mitochondria are integrative hubs central to cellular adaptive pathways. Such pathways are critical in highly differentiated post-mitotic neurons, the plasticity of which sustains brain function. Consequently, defects in mitochondrial dynamics and quality control appear instrumental in neurodegenerative diseases and may also participate in cognitive impairments. To directly test this hypothesis, we analyzed cognitive performances in a mouse mitochondria-based disease model, due to haploinsufficiency in the mitochondrial optic-atrophy-type-1 (OPA1) protein. While in Dominant Optic Atrophy (DOA) models, the known main symptoms are late onset visual deficits, we discovered early impairments in hippocampus-dependent spatial memory attributable to defects in adult neurogenesis. Moreover, less connected hippocampal adult-born neurons showed a decrease in mitochondrial content. Remarkably, modulating mitochondrial function through voluntary exercise or pharmacological treatment restored spatial memory.

Altogether, our study identifies a crucial role for OPA1-dependent mitochondrial functions in adult neurogenesis, and thus in hippocampal-dependent cognitive functions. More generally, our findings show that adult neurogenesis is highly sensitive to mild mitochondrial defects, generating impairments in spatial memory that can be detected at an early stage and counterbalanced by physical exercise and pharmacological targeting of mitochondrial dynamics. Thus, early amplification of mitochondrial function appears beneficial for late-onset neurodegenerative diseases.

## Introduction

Mitochondria are a dynamic population of organelles that migrate, fuse and divide. In doing so, mitochondria adapt to environmental changes at all life stages (Arrazola et al., 2019). As integrative hubs, mitochondria play a particularly important role in highly plastic and fast-responding cells like neurons. The compartmentalization of neurons and their arborization, including spine-endowed dendrites, require not only active and healthy mitochondria, but also tightly regulated mitochondrial transport and dynamics (Flippo & Strack, 2017). Consequently, mitochondrial functions and dynamics exert a pleiotropic influence on neuronal health, from neuronal development to neurodegeneration (Arrazola et al., 2019; Panchal & Tiwari, 2019).

Expectedly, defects in proteins critical for mitochondrial dynamics are directly associated with specific neurodegenerative diseases (Bertholet et al., 2016). Among them, dominant optic atrophy (DOA), a rare disease predominantly affecting the retinal ganglion cells (RGCs) of the optic nerve, is mainly caused by mutations in the gene coding the mitochondrial protein OPA1, involved in inner membrane fusion. In 20% of DOA patients, *OPA1* mutations affect not only RGCs, but also nervous tissues, leading to various neuronal extra-ocular symptoms (Lenaers et al., 2021).

Investigating the role of OPA1 in neuronal functions, we previously demonstrated its critical involvement in neuronal maturation and plasticity (Bertholet et al., 2013). Down-regulation of OPA1 in primary neurons indeed leads to the formation of small, fragmented mitochondria that are unevenly distributed in the dendrites and in which respiration is affected (Bertholet et al., 2013; Millet et al., 2016). These disturbances generate restrictions on dendritic growth, synaptogenesis and a reduction of synaptic proteins (Bertholet et al., 2013). Interestingly, although OPA1 knock-out is embryonically lethal, mouse models carrying heterozygous *OPA1* mutations exhibit late onset RGCs dysfunctions but reach adulthood without reported neuronal developmental problems (Lenaers et al., 2021). Mitochondrial dysfunctions due to OPA1 haploinsufficiency progressively build up, mainly as a pro-oxidative stress, sensitizing neurons to further challenges or insults (Millet et al., 2016).

The role of OPA1 in neuronal maturation, prompted us to explore its influence on the maturation of neurons born in the adult hippocampus. These neurons are indeed submitted to a challenge in order to properly migrate and establish synaptic connections to efficiently integrate the existing neuronal network. We hypothesized that, in a mouse model of OPA1 haploinsufficiency (Alavi et al., 2007), dysfunctional mitochondrial dynamics would affect adult hippocampal neurogenesis, leading to impairment of spatial memory processes that specifically rely on adult-born hippocampal neurons, long before any visual impairment. We also evaluated if such cognitive alterations could be corrected by enhancing brain plasticity *via* mitochondrial targeting.

## Results

### Hippocampal adult neurogenesis is impaired in *OPA1* +/−mice

In the dorsal hippocampus of adult mice, new neurons are generated every day in the dentate gyrus, maturate and migrate to become granule cells. Between 3-5 weeks after their birth, they display a particularly distinct function from their mature state, as they are highly excitable and likely to be recruited into neuronal networks supporting memory. Using immunohistochemistry, we found that OPA1 haploinsufficiency, while sparing cell proliferation (evidenced by Ki67 labeling) and neuronal differentiation (DCX labeling) (Fig. 1A, B), alters the survival of adult-generated hippocampal cells (4-week-old BrdU-labeled) (Fig. 1C). Moreover, adult-generated neurons in *OPA1*+/− mice display impaired dendritic spine density (Fig. 1D, E), reflected by a significant reduction of stubby, thin, and mushroom spine subtypes (Fig. 1F). Thus, under OPA1 deficiency, fewer adult-born cells survive through the critical period, and the surviving adult-born neurons exhibit impaired synaptic connectivity.

**Figure 1.**
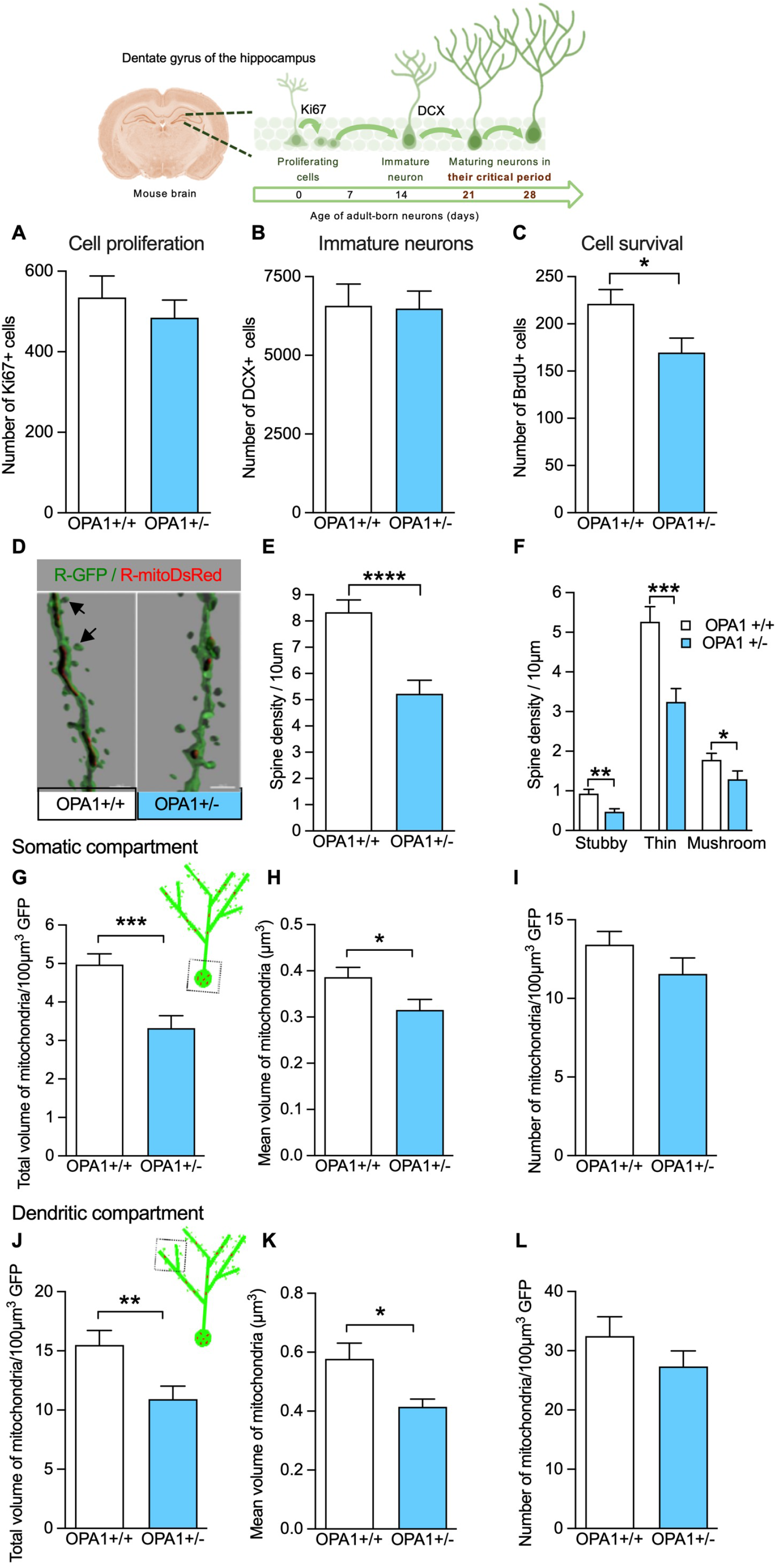
OPA1+/− mice show adult hippocampal neurogenesis impairments. (**A-F**) The status of adult hippocampal neurogenesis in OPA1+/− mice was addressed using immunohistochemistry (7-9 mice / group). Labeling against Ki67 and DCX allowed to evaluate new cell proliferation (**A**) and differentiation towards neuronal fate (**B**), respectively. Numbers of Ki67-labeled (Ki67+) cells (OPA1+/+: 534.8 ± 53.53; OPA1+/−: 484.7 ± 43.57) and DCX+ cells (OPA1+/+: 6575 ± 686.4; OPA1+/−: 6488 ± 558.4) in the dentate gyrus were similar for both genotypes. (**C**) Survival of adult-generated cells was assessed by BrdU labeling. Numbers of 4-week-old BrdU+ cells were significantly lower in OPA1+/− mice compared to controls (OPA1+/−: 221.3 ± 15.09; OPA1+/+: 169 ±1 5.37, *t*=2.39, *df*=13; \**p*<0.05; unpaired t-test). (**D**) Representative 3D reconstruction images showing GFP+ (green) dendritic spines (arrows) and mitoDsRed+ (red) mitochondria. (**E**) In 21-day-old GFP neurons, dendritic spine density was lower in OPA1+/− mice compared to controls (spines/10μm: OPA1+/− 8.34 ± 0.46; OPA1+/+: 5.22 ± 0.51, \*\*\*\**p*<0.0001; Mann-Whitney test). (**F)** Spines were classified based on their morphology. Stubby, thin and mushroom types appeared significantly depleted on OPA1+/− mice GFP-labeled neurons (spines/10μm: *stubby*: OPA1+/+: 0.93 ± 0.10 *vs* OPA1+/− : 0.47 ± 0.08; *thin*: OPA1+/+: 5.27 ± 0.38 *vs* OPA1+/−:3.25 ± 0.30; *mushroom*: OPA1+/+: 1.78 ± 0.16 *vs* OPA1+/−: 1.29 ± 0.21; \**p*<0.05 **, *p*<0.01, \*\*\**p*<0.001; Mann-Whitney test). (**G-L**) Mitochondrial parameters were evaluated in the somatic and dendritic compartments of 21-day-old neurons. (**G, J**) Total mitochondrial biomass was ~30% lower in GFP neurons of OPA1+/− compared to control mice (*somas*: OPA1+/+: 4.98 ± 0.38, OPA1+/−: 3.32 ± 0.33; \*\*\**p*<0.001; *dendrites*: OPA1+/+: 15.49±1.22, OPA1+/−: 10.93±1.08; \*\**p*<0.01, Mann-Whitney test). (**H, K**) The mean volume of individual mitochondria was lower in OPA1+/− compared to control mice (in μm3, *somas*: OPA1+/−: 0.32 ± 0.02; OPA1+/+: 0.39 ± 0.02; *dendrites*: OPA1+/−: 0.41 ± 0.02; OPA1+/+: 0.58 ± 0.05, \**p*<0.05; Mann-Whitney test). (**I, L**) However, the number of mitochondria per somatic or dendritic GFP+ volume was not significantly different between genotypes (particles/100μm^3^, *somas*: OPA1+/+:13.41 ± 0.85, OPA1+/: 11.57 ± 1.01; *dendrites*: OPA1+/+: 32.47 ± 3.28, OPA1+/−: 27.35 ± 2.64). 5 mice/group, 5-7 somas/mouse, 6-8 dendritic segments/mouse.

### Maturing adult-born neurons from *OPA1* +/− mice show altered mitochondrial biomass

In both somatic and dendritic compartments of maturing GFP-labeled neurons of *OPA1*+/− mice, we observed a ~30% reduction of total mitochondrial biomass (Fig. 1G, J), caused by a reduction in the average volume of mitochondria (Fig. 1H, K). The number of mitochondria was not significantly affected (Fig. 1I, L). Thus, OPA1 deficiency results in reduced mitochondrial content in adult-born neurons during their critical developmental period.

### *OPA1* +/− mice have normal general behavior and vision

At the behavioral level, 8-9-month-old *OPA1*+/− mice showed similar anxiety level and locomotor activity than control littermates (Supplementary Fig. 1A, B, C), consistent with a previous report (Caffin et al., 2013). Moreover, these animals displayed normal visual learning and memory in the vision-dependent non-spatial Barnes maze (Supplementary Fig. 2A-D). Similar results were obtained with 4-month old animals (not shown). These findings allowed us to proceed with visually-guided memory tasks.

### *OPA1* +/− mice show spatial memory deficits

Next, we evaluated spatial (hippocampal-dependent) and non-spatial (hippocampal-independent) memory of OPA1 deficient mice. In the spatial version of the Barnes maze (Fig. 2A), *OPA1*+/− mice showed spatial learning and memory (Fig. 2B, C). However, compared to control mice, *OPA1*+/− animals displayed weaker memory robustness (Fig 2D), confirmed at a later delay (Supplementary Fig. 3A, B). Then, these mice were submitted to the object location task, which evaluates the ability of mice to discriminate objects in a novel *vs* a familiar location. This task assesses spatial memory and is sensitive to adult neurogenesis depletion (Goodman et al., 2010). In this task (Fig. 2E), *OPA1*+/− and control littermates showed similar interest for the objects during the exploration phase (Supplementary Fig. 3C). During the test, *OPA1*+/− mice showed no preference for the displaced object, in contrast to control mice (Fig. 2F), revealing impairment of spatial memory. Animals were also tested in the object recognition task, which evaluates non-spatial memory and is insensitive to hippocampal neurogenesis depletion (Goodman et al., 2010). In the non-spatial object recognition task (Fig. 2G), mice of both genotypes exhibited similar interest for both objects during acquisition (Supplementary Fig. 3D). During the test, both genotypes preferentially explored the new object (Fig. 2H). Hence, OPA1 deficiency results in impairment of spatial memory while sparing non-spatial memory.

**Figure 2.**
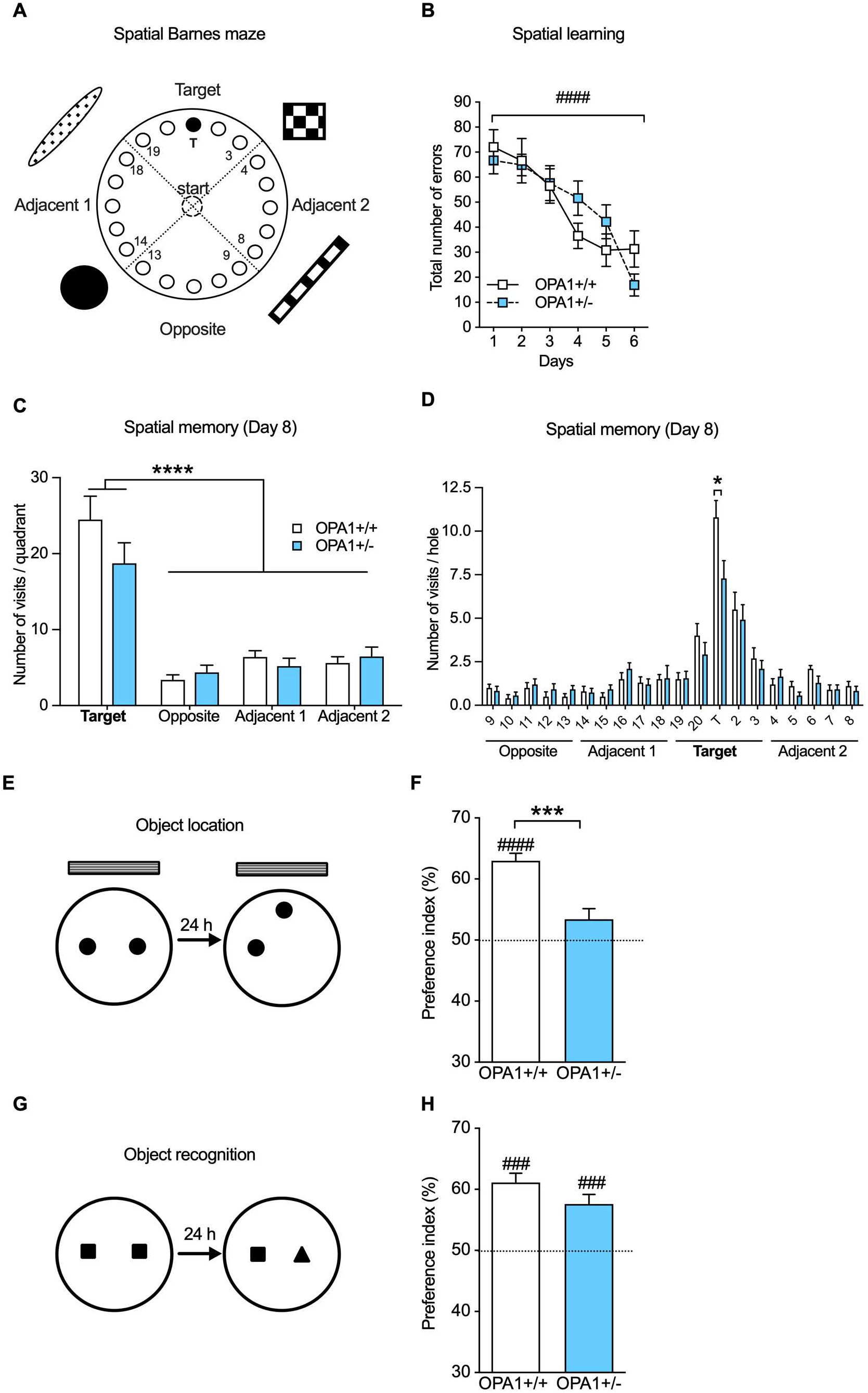
OPA1+/− mice have impaired spatial memory but intact non-spatial memory. (**A**) Schematic setup of the spatial version of the Barnes maze. (**B**) The number of errors before entering into the escape hole decreased across training sessions. OPA1+/+ (*n*=10) and OPA1+/− (*n*=11) mice exhibited similar learning performances during training days (*genotype* : *F*_(1, 19)_=0.025, *p*=0.88; *session*: *F*_(5, 95)_= 25.92, ^####^p<0.0001; two-way ANOVA with repeated measures). Two days after the last training session (Day 8), memory was evaluated by a spatial memory probe test. (**C**) Mice from both genotypes visited significantly more often the holes located in the target quadrant than in the other quadrants (*F*_(3, 76)_=48.8, ****p<0.0001; two-way ANOVA). (**D**) However, OPA1+/− mice visited the target hole significantly less often than the OPA+/+ mice (*t*= 2.502, *df*=19, * *p*<0.05; unpaired t-test). (**E**) Spatial memory was also evaluated in the object location task. (**F**) In contrast to controls, OPA1+/− showed no preference for the displaced object (OPA1+/+, *n*=13: 62.98 ± 1.24%, *t*=10.450, *df*=12, ^####^*p*<0.0001; OPA1+/−, *n*=12: 53.40 ± 1.77%, p>0.05; index *vs* 50%, t-test). Compared to OPA1+/+ mice, mutant mice exhibited a spatial memory deficit (t=4.489, *df*=23, \*\*\**p*<0.001; unpaired t-test). (**G**) Non-spatial memory was assessed in the object recognition task. (**H**) During memory testing, both genotypes explored preferentially the novel object than the familiar one (OPA1+/+, *n*=14: 61.14 ± 1.51%; OPA1+/−, *n*=12: 57.60 ± 1.57%, *t*=7.394, *df*=13 and *t*=4.828, *df*=11, ^###^*p*<0.001 respectively; index *vs* 50%, t-test). In F and G, dotted lines indicate equal exploration of the two objects.

In the metric spatial pattern separation task (Fig. 3A), time spent exploring the objects during exposition decreases significantly over time for mice of both genotypes (Fig. 3B), reflecting habituation to the objects. During the test, *OPA1*+/− mice did not show an increased exploration of the objects in their new metric configuration, unlike control animals (Fig. 3C). Thus, OPA1 haploinsufficiency results in failure to detect changes in the distance between a pair of objects. Overall, our data demonstrate that OPA1 deficiency leads to memory impairment in hippocampal-dependent tasks (spatial navigation in the Barnes maze, object location and pattern separation) while sparing memory in hippocampal-independent tasks (object recognition, cued Barnes maze). This phenotype reveals an important role for OPA1 in hippocampal plasticity. The hippocampal-dependent spatial tasks used here are vulnerable to adult neurogenesis deficits (Clelland et al., 2009; Goodman et al., 2010; Raber et al., 2004), which led us to ask whether stimulation of adult hippocampal neurogenesis and/or the mitochondrial system could reduce/abolish the observed cognitive defects.

**Figure 3.**
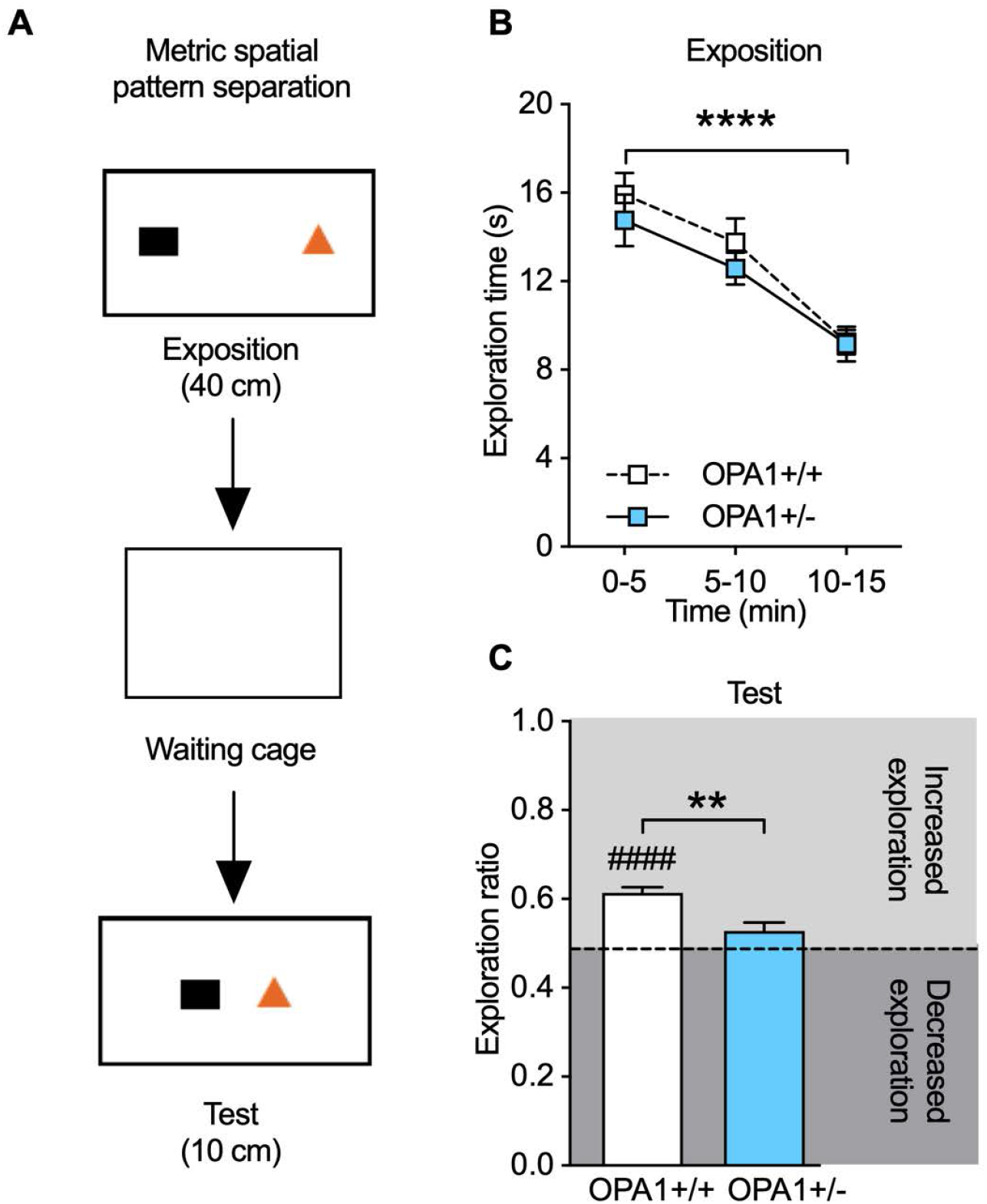
OPA1+/− mice show deficits in metric spatial pattern separation. (**A**) Schematic of the pattern separation procedure. (**B**) For both genotypes, habituation to the objects was reflected by a decrease in exploration time during exposition phase (*session*: *F*_(1.407, 30.95)_=25.36, \*\*\*\**p*<0.0001; two-way ANOVA with repeated measures). No genotype effect was observed (*F*_(1,22)_=0.994, *p*=0.33). (**C**) During memory test, the exploration ratio increased significantly in OPA1+/+ mice (OPA1+/+, *n*=12: 0.61 ± 0.01; t=8.914, *df*=11, ^####^ *p*<0.0001) while the exploration ratio of OPA1+/− was not different from 0.5 (OPA1+/−, *n*=12: 0.53 ± 0.02; *t*=1.430, *df*=11, p=0.181; t-test). Performances in pattern separation were significantly different between genotypes (*t*=3.675, *df*=22, \*\**p*<0.01; unpaired t-test).

### Voluntary exercise rescues spatial memory deficits in *OPA1* +/− mice

First, we investigated the consequences of voluntary exercise, known to enhance adult neurogenesis (Van Praag, Kempermann, & Gage, 1999), on spatial memory deficits in *OPA1*+/− mice (Fig. 4A). When housed under standard conditions, *OPA1*+/− mice showed no preference for the displaced object, revealing a spatial memory impairment, in contrast to their *OPA1*+/+ counterparts (Fig. 4B). However, after a 3-week period of voluntary running, *OPA1*+/− mice preferentially explored the displaced object (Fig. 4B), showing ability to distinguish the new location from the familiar one. Thus, voluntary exercise abolishes the spatial memory deficit of OPA1-deficient mice. Consistent with this observation, running also results in a significant (33%) increase in somatic mitochondrial content of adult-born neurons in *OPA1*+/− mice compared to *OPA1*+/− housed in standard conditions (total mitochondrial volume / 100μm^3^ of GFP in the soma: 5,072 ± 0,475 *vs* 3,79 ± 0,349, p<0,038, Mann-Whitney test; data not illustrated).

**Figure 4:**
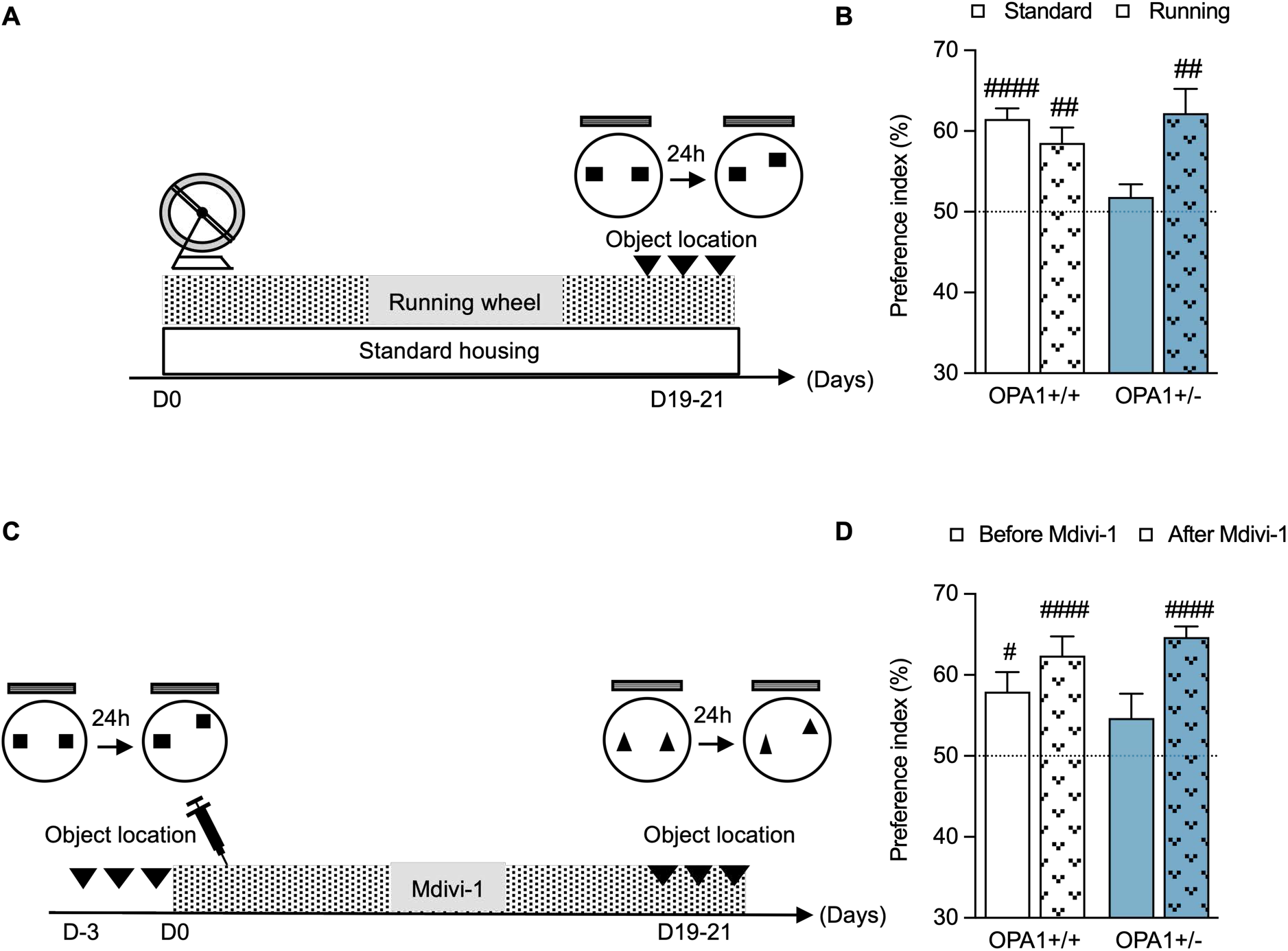
Both voluntary running and Mdivi-1 treatment rescue hippocampal-dependent memory deficits in OPA1+/− mice. **(A)** Mice had access to voluntary running activity or remained in standard conditions for 3 weeks before being tested in the object location task. (**B**) OPA1+/− mice enrolled in physical exercise explored preferentially the displaced object (OPA1+/− run, *n*=7: 62.24 ± 2.99; *t*=4.096, *df*=6, *p*<0.01), in contrast to OPA1+/− mice housed in standard conditions (OPA1+/− standard, *n*=8: 51.85 ± 1.56, *t*=1.184, *df*=7, *p*=0.275; index *vs* 50%, t-test). OPA1+/+ mice housed in both standard and running conditions discriminated the displaced object (OPA1+/+ standard, *n*=8: 61.50 ± 1.30, *t*=8.849, *df*=7, ^###^*p*<0.0001; OPA1+/+ run, *n*=7: 58.54 ± 1.91, *t*=4.452, *df*=6,^##^*p*<0.01, respectively), indicating that voluntary running abolished the impairment of object location performance in OPA1-deficient mice. (**C**) Mice were tested for spatial memory before and after 3 weeks of Mdivi-1 treatment. (**D**) After Mdivi-1, OPA1+/− mice preferentially explored the displaced object (*n*=9: 64.67 ± 1.35; *t*=10.90, *df*=8, ^####^*p*<0.0001) unlike OPA1+/− mice before treatment (54.67 ± 3.01; *t*=1.554, *df*=8, *p*=0.158; index *vs* 50%, t-test). OPA1+/+ mice explored preferentially the displaced object, both before and after Mdivi-1 (*n*=9: 57.94 ± 2.41, *t*=3.298, *df*=8, ^#^*p*<0.05; *n*=9: 62.37±2.41, *t*=5.138, *df*=8, ^####^*p*<0.0001).

### Pharmacological stimulation of mitochondria rescues spatial memory in *OPA1* +/− mice

In order to pharmacologically target mitochondria, we used Mdivi-1, an inhibitor of the DRP1 mitochondrial fission protein (Cassidy-Stone et al., 2008; Rosdah, J, Delbridge, Dusting, & Lim, 2016), known to compensate for the lack of fusion related to OPA1-deficiency. Mice from both genotypes were tested in the object location task before and after Mdivi-1 treatment (Fig. 4C). During the exploration phase, all mice spent the same amount of time exploring each object (data not shown). Before treatment, *OPA1*+/− mice showed no preference for the displaced object, in contrast to *OPA1*+/+ littermates (Fig. 4D). Remarkably, after treatment, *OPA1*+/− mice significantly explored the displaced object (Fig. 4D), indicating that they were now able to discriminate the new location from the familiar one. Consequently, treatment with Mdivi-1 rescued spatial memory of *OPA1*+/− mice.

Thus, both physical exercise increasing mitochondrial content in adult-born neurons (Steib, Schaffner, Jagasia, Ebert, & Lie, 2014) and pharmacological treatment targeting mitochondrial dynamics correct hippocampal-dependent memory deficits in *OPA1*+/− mice. As performance in these tasks is highly dependent on adult neurogenesis (Clelland et al., 2009; Goodman et al., 2010; Sahay et al., 2011), our data suggest that recruitment of adult-generated neurons into hippocampal neural networks supporting this type of memory is particularly vulnerable to mitochondrial dysfunction and can be restored by acting on mitochondria.

## Discussion

Our data show for the first time that maturing adult-born hippocampal neurons are particularly vulnerable to the alteration of mitochondrial dynamics induced by a reduction of the mitochondrial inner membrane protein OPA1 in mice. In these *OPA1*+/− mice, the restricted number of new neurons and their limited connectivity within the existing network lead to cognitive impairments. Furthermore, we provide evidence that indirectly acting on the mitochondrial content alleviates such cognitive deficits in OPA1-happloinsufficient mice.

Stem cell proliferation and differentiation depend on controlled events including regulation of mitochondrial dynamics (Arrazola et al., 2019; Giacomello, Pyakurel, Glytsou, & Scorrano, 2020). Beyond the well-known shift from glycolysis to oxidative phosphorylation, pioneer gain- and loss-of-function studies targeting key actors of mitochondrial functions and dynamics showed that distinct metabolic states are critical for each developmental step of neurons born during adulthood in the hippocampus, particularly during early lineage progression (Beckervordersandforth et al., 2017; Khacho et al., 2017). In line with these data, we find that OPA1 haploinsufficiency leads to impairments in the survival, connectivity, and mitochondrial content of adult-born granule neurons. However, the restricted OPA1 content in the *OPA1*+/− mouse model impacts neither proliferation nor neuronal fate of neural stem cells in the hippocampus. Nevertheless, newborn cells display reduced survival, suggesting a lack of pro-survival events during this period. Neurons generated during adulthood go through a “critical period” of maturation wherein dendritic arborization and spinogenesis take place, two processes requiring proper mitochondrial functions and dynamics (Arrazola et al., 2019). In line with this, OPA1 deficiency is associated with defects in spine density as well as mitochondrial content of adult-born granule neurons. Specifically, mitochondria are smaller, more sparsely distributed along dendrites, and thus may be less able to support spinogenesis and the high neuronal activity supporting memory processes during their critical period.

Among spine subtypes, stubby are considered immature, whereas thin and mushroom ones are known to be associated with active synapses (Nimchinsky, Sabatini, & Svoboda, 2002). All these subtypes show reduced density in *OPA1*+/− mice, further documenting the intricate interplay between mitochondrial dynamics and synaptic connectivity in the mouse brain. In maturing primary neurons, mitochondrial defects due to OPA1 down-regulation lead to impaired spinogenesis (Bertholet et al., 2013). Reciprocally, synaptic activity regulates the number of spine-associated mitochondria (Rangaraju, Lauterbach, & Schuman, 2019; Seager, Lee, Henley, & Wilkinson, 2020). In the adult hippocampus, maturing new neurons display highly plastic properties that favor their functional integration (Jessberger & Kempermann, 2003). Our findings thus show that OPA1 reduction impinges on the critical period of maturation of adult-born neurons and suggest that synaptic activity might be limited in the dentate gyrus of *OPA1*+/− mice.

Consistent with the decrease in spine density of adult-born neurons observed in *OPA1*+/− mice, we provide the first evidence of spatial memory defects in these mice at middle age (8-9 months of age), attributable to cognitive deficits. Their visual capacity is indeed maintained, as shown using conventional vision-based tests (Pinto & Enroth-Cugell, 2000), and these mice display intact RGCs number and optic nerve myelination at 11 months of age (Atamena et al., unpublished results). In agreement, these animals, like most DOA patients, display late-onset visual deficits due to optic nerve degeneration (Lenaers et al., 2021). Using another DOA mouse model, a recent study reported cognitive impairments at older age (over 14 months), when visual impairments are described, precluding reliable determination of the cognitive profile of these mice at this age (Bevan et al., 2020).

Here, we demonstrate that these animals display normal locomotor activity, motivation, and anxiety level. They also show intact performances in cued (visual non-spatial) memory test and in hippocampal-independent recognition test at middle age.

In the hippocampus, adult-born neurons within their critical period of maturation and integration are repeatedly shown to play a crucial role in memory processes related to spatial navigation and pattern separation (Goodman et al., 2010; Sahay et al., 2011; Trouche, Bontempi, Roullet, & Rampon, 2009). Hence, we associate the cognitive impairments in *OPA1*+/− mice evidenced here in object location and metric-spatial pattern separation tasks with defects in these maturing neurons. Since voluntary exercise is known to enhance adult neurogenesis (Van Praag et al., 1999) and to potentiate the reciprocal interactions between mitochondria and maturation of adult-born neurons (Steib et al., 2014), the rescuing effect that we observe on spatial memory and likely on new neurons mitochondrial biomass in *OPA1*+/− mice further suggests a role for mitochondria in these defects. More specifically, mitochondrial dynamics appears crucial, as administration of Mdivi-1, a pharmacological inhibitor of the mitochondrial fission protein DRP1 (Cassidy-Stone et al., 2008; Rosdah et al., 2016), counteracted spatial memory deficits in *OPA1*+/− mice. Whether or not the *in vivo* effects of Mdivi-1 treatment are direct or indirect, or related to other mitochondrial parameters than DRP1 inhibition, as suggested by some recent reports (Aishwarya et al., 2020), remains to be investigated. However, Mdivi-1 treatment was repeatedly proven to be neuroprotective, *in vitro* against axonal damage induced by rotenone (Lassus et al., 2016) as well as *in vivo* in Alzheimer’s disease context (Wang et al., 2017). This points to mitochondrial dynamics as a therapeutic target.

### Overall, demonstrating the reversibility of the consequences of OPA1 haploinsufficiency on spatial memory highlights the key involvement of mitochondrial dynamics in the unique plasticity of adult-born neurons

Mitochondria-based defects could therefore be revealed at an early stage of the DOA pathology by spatial memory tests, calling for further investigation in patients with DOA. Finally, the beneficial effects of pharmacological targeting of mitochondrial dynamics on adult neurogenesis-dependent cognitive processes open new therapeutic possibilities for neurodegenerative diseases.

## Materials and methods

Detailed methods are available in the Supplementary material.

### Animals

We produced *OPA1*329-355del (*OPA1*+/−) mice (from Alavi et al, 2007, kind gift from Dr B. Wissinger, University of Tubingen) in our animal facility, PCR-genotyped on *Opa1* gene exon 10 (Alavi et al., 2007). Experiments were conducted on males.

### Ethics statement

All experiments were performed in strict accordance with the policies of the European Union (2010/63/EU) for the care and use of laboratory animals. Animal facility of the CRCA is fully accredited by the French Direction of Veterinary Services (D 31–555–11, Sep 19, 2016) and experimental procedures conducted in this study were authorized by local ethical committees and the French Ministry for Research (#12342-2017082111489451 v6). All efforts were made to improve animal welfare and minimize animal suffering.

### Histological experiments

#### Stereotactic retroviral injections

We used MML-retroviral vectors specifically transducing proliferating cells and previously described (Roybon et al., 2009); R-GFP (pCMMP-IRES2-eGFP-WPRE) for cytoplasmic expression of eGFP (kind gift of Dr. L Roybon, Lund University) and R-mitoDsRed (pCMMP-IRES2-mitochondrial Discosoma Red-WPRE) for expression of mitochondrial matrix-targeted DsRed (Steib et al., 2014) (kind gift of Dr. DC Lie, Helmholtz Zentrum München), at final titers of 0.7-1 x10E9 TU/ml. Mice were anaesthetized (4% isoflurane), maintained under 2.5-3% Isoflurane, and bilaterally injected (1 μL of a 1:1 retrovirus mix) into the dentate gyri (Richetin et al., 2017).

#### BrdU and Mdivi-1 administration

Mice received three intraperitoneal injections (100mg/kg) of 5’-bromo-2 deoxyuridine (BrdU, Sigma) at 4 h intervals and were sacrificed after 28 days. Mdivi-1, 3-(2,4-Dichloro-5-methoxyphenyl)-2-sulfanyl-4(3H)-quinazolinone (BML-CM127-0050, Enzo Life Sciences) was injected at 20mg/kg (in 10% DMSO) intraperitoneally, three times a week for three weeks.

#### Tissue processing and immunohistochemistry

Anaesthetized mice were transcardially perfused. Brains were post-fixed, cryoprotected, sectioned and used for immunostainings as previously described (Richetin et al., 2017; Verret, Jankowsky, Xu, Borchelt, & Rampon, 2007).

#### Cellular stereological counting

Stereological quantification of BrdU, Ki67 and DCX immunolabeled-cells was conducted using Mercator software (Explora Nova, France).

#### Confocal analysis of dendritic spines and mitochondrial content

GFP+ neurons expressing mitoDsRed were imaged with a confocal microscope. Images were deconvoluted (Huygens Essential deconvolution) and analyzed using the 3D Imaris XT software (Bitplane AG). Dendritic spine density and morphology were analyzed using GFP staining. Mitochondrial parameters were determined for 100μm^3^ of GFP volume.

### Behavioral experiments

Behavioral tests were performed by experimenter blind to genotypes, using published protocols. *OPA1*+/+ and *OPA1*+/− mice were tested for anxiety-related and exploratory behaviors. The non-spatial (cue) version of the Barnes maze (vision-guided) and the object recognition task allowed to evaluate hippocampal-independent memory. The spatial version of the Barnes maze, the pattern separation and object location tasks were used to evaluate hippocampal-dependent memory. Consequences of voluntary exercise were assessed using a standard running procedure. All parameters were measured using Ethovision software (Noldus, The Netherlands).

### Statistics

Statistical analyses were performed using Prism software (GraphPad.9). Appropriate analyses are indicated in the legends of the figures. All data are presented as mean ± SEM.

### Data availability

The data that support the findings of this study are available from the corresponding author, upon reasonable request.

## Abbreviations

BrdU: 5’-Bromo-2 deoxyuridine
DCX: Doublecortine
DAB: di-amino-Benzidine
DMSO: Dimethyl sulfoxide
DOA: Dominant optic atrophy
Mdivi-1: Mitochondrial division inhibitor 1
PB: Phosphate buffer
PBST: Phosphate-buffered saline with Tween 20
PFA: Paraformaldehyde
OPA1: Optic atrophy type 1 protein
RGCs: Retinal ganglion cells

## Acknowledgements

The authors thank Vincent Setola, Laure Verret, Ambre Bertholet, Lucienne Ronco, Nicholas Rhind, Jean-Michel Peyrin and Didier Vilette-Miquel for critically reading the article, Aude Deleau and Stéphane Pech for animal care and technical help. Mice were housed in the ABC Facility of ANEXPLO, Toulouse, behavioral testing was performed on the CBI-Mouse Behavioral Core (MBC), confocal analysis was run on the CBI-LITC platform.

## Supplementary material

Supplementary material is available at bioRxiv online.

