## Supplemental figures and legends for "Mitochondrial OPA1 deficiency causes reversible defects in adult neurogenesis-associated spatial memory in mice"

### Supplementary Figure & legends

#### Supplementary Figure 1

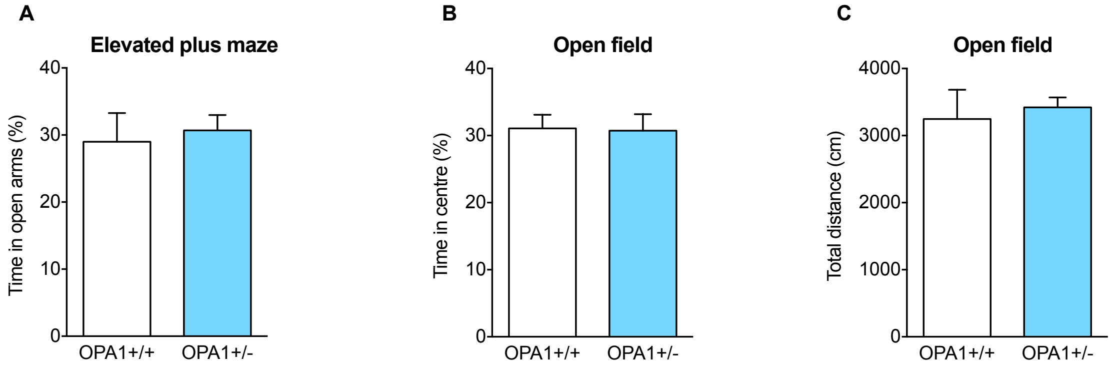


**Supplementary Figure 1**

**Intact anxiety-related and locomotor behavior in *OPA1*+/- mice**

OPA1 deficiency has no impact on anxiety-like behavior measured by **(A)** the time spent in the open arms of the elevated plus maze (in %, *OPA1*+/+: 29.01±4.27; *OPA1*+/-: 30.72±2.28, $t$=0.34, $df$=25, $p$=0.734; t-test). Mice from both genotypes show similar locomotor activity in the open-field, evaluated by **(B)** the time spent in the center area (in %, *OPA1*+/+:31.07±2.03; *OPA1*+/-: 30.73±2.46, $t$=0.11, $df$=25, $p$=0.915; t-test), and (**C**) the total distance travelled (in cm, *OPA1*+/+ = 3246 ± 437.7; *OPA1*+/- = 3423 ± 145.1, $p$=0.239; Mann-Whitney test). *OPA1*+/+, $n$=14; *OPA1*+/-, $n$=13.

### Supplementary Figure 2

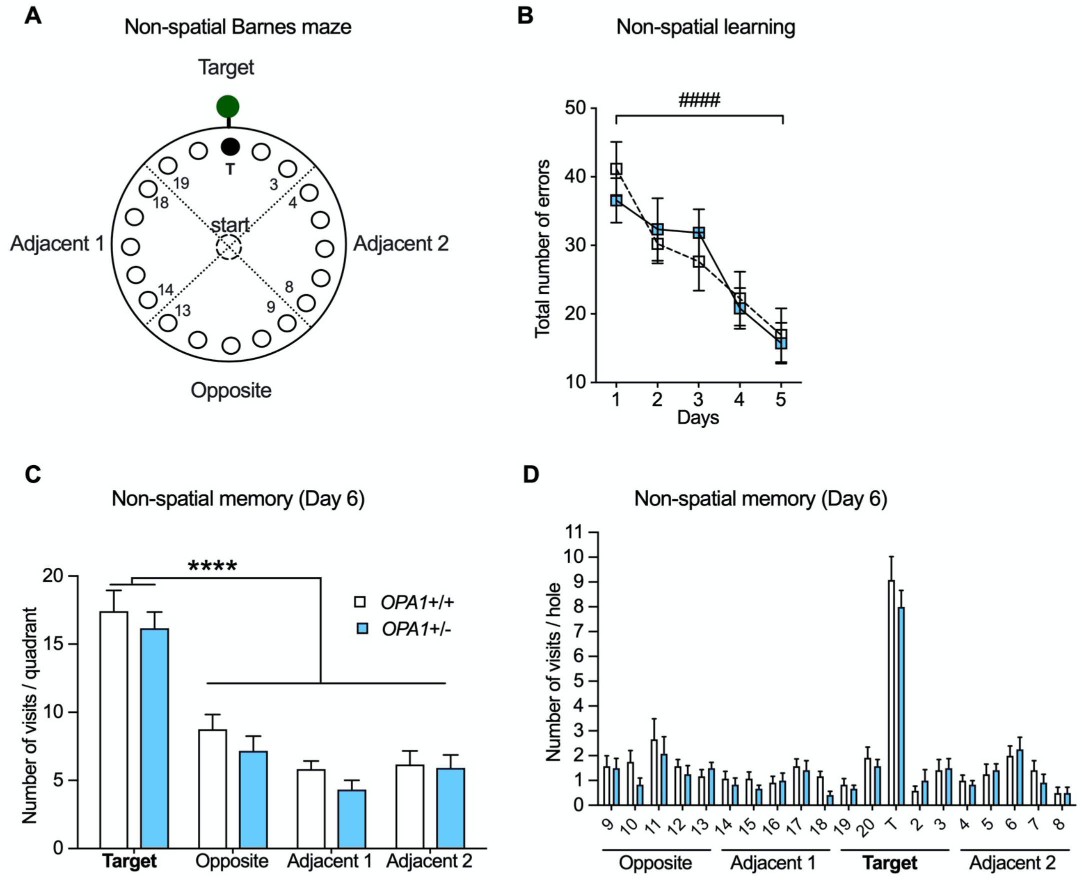


**Supplementary Figure 2**

***OPA1*+/- mice display intact performances in the non-spatial Barnes maze**

**(A)** Schematic setup of the non-spatial (cued) Barnes maze. (**B**) During the five days of training, the number of visits before entering into the escape hole decreased for mice of both genotypes, indicating they learned to locate the escape hole (*session*: $F_{(4,88)}$=17.82, $^{\#\#\#\#}p$<0.0001; two-way ANOVA with repeated measures). No difference was found between genotypes ($F_{(1,22)}$=0.002, $p$=0.96). (**C**) During the non-spatial probe test held 24 hours after training (Day 6), both groups of mice visited more often the holes located in the target quadrant than in each other quadrants ($F_{(3,\ 88)}$=51.35, ****$p$<0.0001) and no difference was observed between genotypes ($F_{(1,88)}$=2.355, p=0.128). (**D**) *OPA1*+/+ and *OPA1*+/- mice exhibited a similar preference for the target hole ($p$=0.3596). *OPA1*+/+, $n$=12; *OPA1*+/-, $n$=12.

Supplementary Figure 3

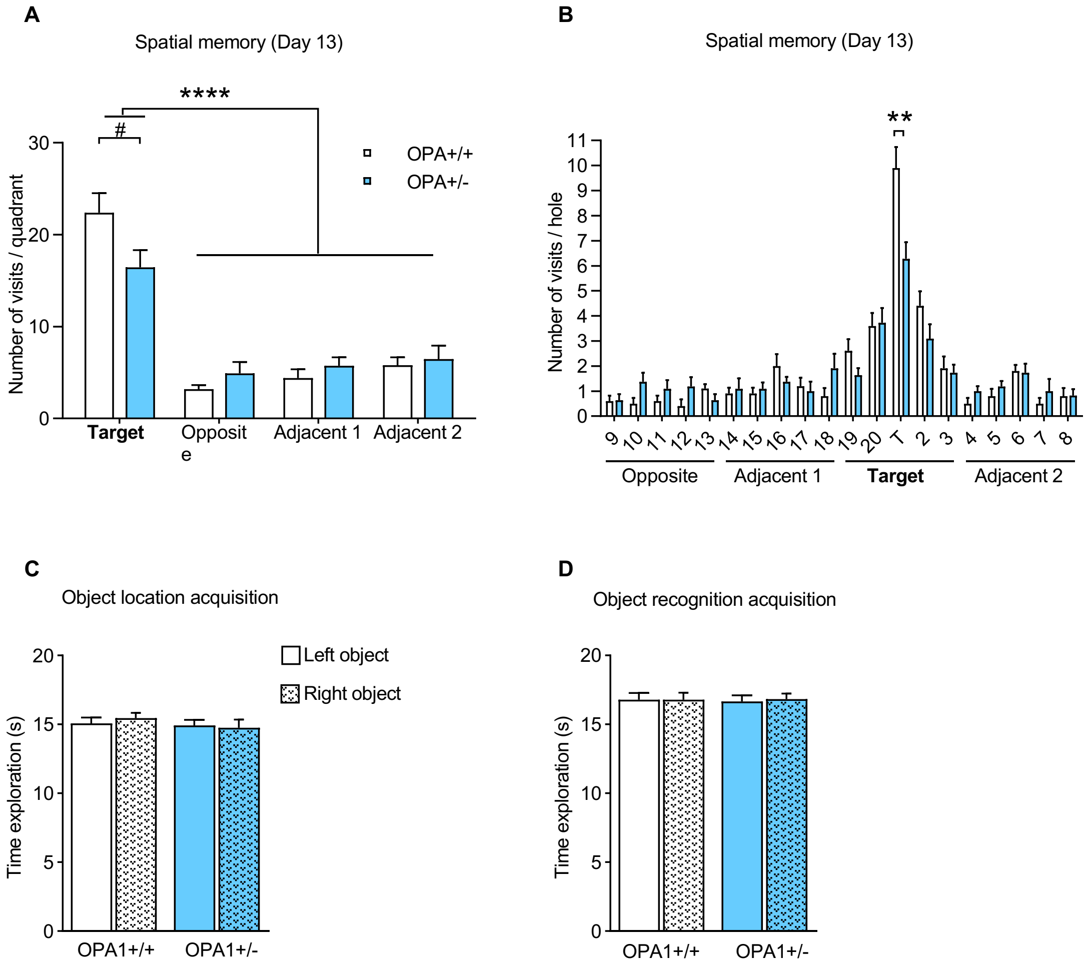


**Supplementary Figure 3**

**Long-term spatial memory is less accurate in *OPA1*+/- mice than in *OPA1*+/+ mice**

(**A**) During the spatial memory probe test held 7 days after training (Day 13), mice from both genotypes visited significantly more often the holes located in the target quadrant than those located in the other quadrants ($F_{(3, 16)} = 48.8$, ****$p<0.0001$; two-way ANOVA). Moreover, *OPA1*+/- visited significantly less often the holes of the target quadrant ($^{\#}p<0.05$, Bonferroni's multiple comparison). (**B**) This effect was strictly attributable to a difference between *OPA1*+/- and *OPA1*+/+ mice in the number of visits to the target hole ($t$=3.433, $df$=19, ** $p<0.01$; unpaired t-test), revealing a less precise spatial memory in mutant mice. *OPA1*+/+, *n*=10; *OPA1*+/-, *n*=11). ***OPA1*+/+ and *OPA1*+/- mice express similar interest for all objects.** Mice from both genotypes spent the same amount of time exploring the left and right objects during the acquisition phase of the object location **(C)** and object recognition **(D)** tasks. Mean exploration time was not different between genotypes, indicating that both groups of mice had a similar interest for each object. *Object location*: *OPA1*+/+, *n*= 13; *OPA1*+/-, *n*=12; *Object recognition*: *OPA1*+/+, *n*= 14; *OPA1*+/-, *n*=12.
