## Supplementary information Materials & methods for "Mitochondrial OPA1 deficiency causes reversible defects in adult neurogenesis-associated spatial memory in mice"

#### Supplementary methods

##### Animals

Mice from the heterozygous OPA1329-355del (OPA1+/-) strain developed by Alavi[1] et al, (2007) were generated in our animal facility (ABC, Anexplo, Toulouse) by crossing OPA1+/- mice with C57BL6JRj females (Janvier Labs, France) to obtain OPA1+/- and OPA1+/+ control littermates. Genotyping was done by PCR analysis of the presence of the exon 10 of the *Opa1* gene, as previously described.[1]-Splice site mutation in the *Opa1* gene of OPA1+/- mice leads to skipping of exon 10 and consequently, OPA1+/- mice show a 50% reduction in OPA1 protein levels compared to OPA1+/+ mice.[1] Males of 4 or 8-9 months of age were used for the anatomical and behavioral studies, respectively.

##### Housing conditions

In standard conditions, mice were maintained by 4-5 animals per standard cage, under a 12/12 h light/dark cycle, with free access to food and water. In running conditions, mice were housed in larger cages (425×266×185 mm) containing running wheels (2 wheels/ 4 mice) to which animals had free access during 3 weeks.

##### Injection of retroviral vectors through stereotaxic surgery

Enhanced green fluorescence protein (GFP) and red fluorescent mitoDsRed Moloney murine Leukemia-derived retroviral vectors (pCMMP-MCS-IRES-eGFP and pCMMP-IRES2-mDsRed-WPRE) were used to label hippocampal adult-born neurons and their mitochondrial content. These vectors were kindly produced by Dr. Roybon (Lund University, Sweden).

Mice were anesthetized with 4% isoflurane and placed in a stereotaxic apparatus (Stoelting) with a mask connected to maintain the anesthetic with 2.5-3% Isoflurane during the entire surgery. Each mouse was bilaterally injected with 1 µl of a 1:1 retrovirus mix (0.1 ml/min), into the dentate gyrus (relative to bregma: anteroposterior -2 mm; lateral ±1.6 mm; ventral -2.5 mm; as in Krezymon[2] et al., (2013). After injections, lidocaine was applied on the flesh before suturing the skin and the mice were placed under a heating lamp until recovery, then returned to home cage. Mice were sacrificed 3 weeks after viral injections, in order to evaluate the morphology and mitochondrial content of 21-day-old new granule neurons after they have begun their synaptic integration.[3]

### BrdU injections

Mice received 3 intraperitoneal injections of 100 mg/kg of 5'-bromo-2 deoxyuridine (BrdU, Sigma, St. Louis, MO) dissolved in 0.9% NaCl, pH=7.4, at 4 h intervals and were sacrificed 28 days later, in order to evaluate survival of newly born cells.

### Tissue processing

Mice were deeply anesthetized with dolethal solution (2g /kg) (28 days after BrdU injection, or 21 days after viral injection), and transcardially perfused with 0.1 M phosphate buffer (PB 0.1M) followed by 4% paraformaldehyde (PFA). Brains were removed and fixed overnight in 4% PFA, rinsed 24 hours in PBS and cryoprotected in a 30% sucrose solution containing 0.1% sodium azide, at 4°C for at least 2 days. Brains were cut into 40 µm thick coronal sections using a sliding microtome (Leica SM2010R) equipped with a freezing-stage (Physitemp BFS-3MP). Sections were kept in cryoprotectant solution at -20°C until use.

### BrdU, Ki67 and DCX immunohistochemistry

Free-floating brain sections were rinsed overnight in 0.1M PBS with 0.25% Triton X-100 (PBST) before being submitted to immunohistochemistry directed against Ki67 (an endogenous marker of proliferating cells), DCX (a marker of immature neurons) or BrdU (marker birth-dating dividing cells). All steps were run under gentle agitation at room temperature, with PBST rinses (2x20 min) after each incubation. After rinses, sections were processed for quenching of endogenous peroxidases with 10% $H_2O_2$ in 10% methanol in PBS for 15 min. For BrdU staining, sections were incubated in 2N HCl for 50 min to denature DNA followed by neutralization in 0.1M borate buffer (pH 8.5). For all staining, non-specific labeling was prevented by incubation in 5% normal goat serum (NGS) in PBST for 1 hour. Sections were then incubated overnight in either one of the following primary antibody solutions: PBST-5% NGS with rat anti-BrdU (1:400; OBT-0030, Harlan Seralab, Loughborough, UK) or PBST containing 5% NGS, 1% BSA and 0.5% Tween-20 with rabbit anti-human Ki67 (1:500 in; SP6 Mab, Spring Bioscience Corp, USA), or goat anti-DCX (1:100 in PBST; Santa Cruz Biotechnology, USA). The next day, sections were incubated for 90 min at room temperature in one of the following biotinylated secondary antibody solutions: goat anti-rat (1:400), goat anti-rabbit (1:500), or rabbit anti-goat (1:500) (all from Vector Laboratories, USA) in PBST. All sections were then incubated for 90 minutes in avidin-biotin-peroxidase complex (1:400; Vector Laboratories Elite Kit) in PBST. Staining was visualized with 0.05 M Tris-HCl buffer, pH 7.6, containing 0.025% Di-Amino-Benzidine (DAB), 0.003% $H_2O_2$ and 0.06% nickel ammonium sulfate. Reaction was stopped by extensive rinsing with PBST containing 0.1%

sodium azide (PBST-Az). Sections were mounted onto slides, counterstained with Nuclear Fast Red (Vector Laboratories), dehydrated through graded alcohols, and cover-slipped.

#### GFP immunofluorescence staining

After extensive rinsing in PBST, free-floating brain sections from retro-virus injected mice were incubated in a solution of rabbit anti-GFP (1:500, Torey Pines Biolabs) diluted in PBST overnight. Then sections were rinsed in PBST and incubated for 120 min in a solution of Alexa Fluor-488-conjugated donkey anti-rabbit (1:500; Life Technologies) in PBST. Sections were rinsed in PBST, mounted in Mowiol containing Hoechst (1:10 000; Life Technologies) and cover-slipped.

#### Quantification of BrdU, Ki67 and DCX -labeled cells

Stereological quantification of BrdU, Ki67 or DCX -labeled (BrdU+, Ki67+ or DCX+) cells was conducted bilaterally from a 1-in-6 series of sections (240 µm interval) for BrdU staining or from a 1-in-6 series of sections (480 µm interval) for Ki67 or DCX staining, through the rostro-caudal extent of the dorsal hippocampus. Immunolabeled cells were counted manually at 40x magnification using a microscope (Leica DM6000 B) equipped with digital camera (ProgRes CFCool, Jenoptik, Jena). The experimenter was blind to the experimental groups. The density of labeled cells was calculated by dividing the number of BrdU+, Ki67+ or DCX+ cells by the granule cell layer / subgranular zone sectional volume measured with the Mercator morphometric system (Explora Nova, France). The total numbers of labeled-cells were obtained by multiplying these densities by the reference volume, as previously described.[4]

#### Confocal imaging of neuronal morphology and mitochondrial content

To analyze mitochondria and dendritic spines, images of GFP+ neurons expressing mitoDsRed in the dentate gyrus were captured using a confocal microscope (Leica TCS SP8, Germany) equipped with 63x HCX PL APO (1.40 NA) objective. All images were taken in z-series at a 0.20-mm interval, with a 63x oil lens and digital zoom of 6. The same settings for laser power, photomultiplier gain and offset were maintained for all images. The images were acquired with a resolution of 1024 X 1024 pixels, 4 times line and frame averaged and 8000 Hz scanning speed. The image stacks of 12-bit files were then deconvoluted using the Huygens Essential deconvolution software (SVI). All images were imported into the 3D Imaris XT software (Bitplane AG) for dendritic spines and mitochondria content analysis.

#### Analysis of dendritic spine density and shape

Spines were analyzed within the distal dendritic part of adult hippocampal new neurons in order to describe the morphological synaptic integration of these neurons. The distal dendritic compartment was defined as the dendritic shaft located between the beginning of the middle molecular layer and the end of the outer molecular layer, comprising mainly segments located after the second dendritic branching point. Spine analysis included spine density (number of spines/10µm dendritic length) and morphological classification. The 3D reconstruction of the dendrites was performed with the Imaris software. Spines volume and number were determined with the Imaris surface tool module, while dendritic length was measured with the Imaris Filament tracer module. Dendritic spines were defined as protrusions from the dendritic shaft and were categorized based on their individual volume and morphology into 3 types, excluding filopodia: *thin* (protrusion with a neck and a head < 0.1 $\mu m^3$), *stubby* (protrusion with no obvious neck or head) or *mushroom* (protrusion with a neck and a head >0.1 $\mu m^3$).[5] Spine density was calculated by dividing the total number of spines by the length of the dendritic segment, as previously described.[6] Confocal imaging and data analysis were performed blindly to experimental conditions.

#### Mitochondrial content of adult-born hippocampal neurons

Mitochondria inside GFP+ neurons were analyzed in the somatic and distal (as above defined) dendritic compartments. The total volume of mitochondria, their mean volume and their number were automatically counted using the Imaris software (Bitplane). For both the somatic and distal dendritic compartments, mitochondrial total volume was determined for 100$\mu m^3$ of GFP volume of the corresponding compartment, mitochondrial number and mitochondrial mean volumes were determined within the same portions, as previously described.[7] Confocal imaging and data analysis were performed blindly to the experimental condition.

### Behavioral experiments

A comprehensive battery of tests was deployed to evaluate several aspects of mice behavior. Before each experimental session, mice received 3 to 4 days of handling to get habituated to the experimenter. Mice were pseudo-randomly distributed into batches of 8-14 mice with counterbalanced numbers of OPA1+/+ and OPA1+/- genotypes, experimenters being blind to genotypes. All experiments were video-recorded and locomotor activity was automatically analyzed using Ethovision XT software (Noldus, The Netherlands) except tasks involving objects, for which the exploration time was recorded manually. The setups were cleaned thoroughly with 70% ethanol followed by water between each mouse session, to ensure the absence of olfactory cues.

*General behavior:* Anxiety-related behavior was assessed using an elevated plus maze. Locomotor activity was recorded during the 10 min familiarization phase of the object location or recognition tasks, using an empty circular arena (40 cm diameter) devoid of visual cues.

*Spatial and cued navigation in the Barnes maze:* This behavioral paradigm relies on the innate preference of rodents to escape bright light and open spaces.[8, 9] The circular Barnes maze (90 cm diameter) contains 20 holes (5 cm diameter, 8 cm apart) evenly located around the perimeter. It is elevated 90 cm from the floor and receives approximately 800 lux from a centered bright light, in order to motivate the mice to escape through the escape cage. The target hole is connected with a tube to an escape cage located beneath the maze. Other cages containing bedding mix from the escape cage, but not connected to any of the holes, were positioned underneath the maze to avoid olfactory bias. We used two paradigms, and independent groups of mice, to respectively evaluate hippocampal-independent non-spatial and hippocampal-dependent spatial navigation. For the non-spatial (cued, hippocampal-independent) version, a ball (10 cm diameter) was placed near the target hole, and the position of the cued target hole varied pseudo-randomly over days (Supplementary Fig. 3A). Here, mice learnt the association between the cue (the ball) and the target hole. For the spatial version of the task (hippocampal-dependent), visual cues were positioned on the walls surrounding the maze (Fig. 2A). The location of the target hole remained the same across trials and days, so that the mice had to learn the location of the target hole based on the distal cues.

We used a modified protocol of Sunyer[10] et al (2007). Briefly, after 3 consecutive days of handling, mice were submitted to either the cued or spatial protocol and exploration was manually recorded. First, mice were familiarized to the setup and to descending into the escape cage through the tube. Then animals received four trials per day during five (cued) or six (spatial) days. For each trial, the mouse was released from the center of the maze and given 5 min to enter the target hole, after which, in case of failure, the mouse was gently guided to the target hole. For each trial, the total number of errors made before first entrance into the target hole was recorded. Errors are scored each time a mouse visits (i.e., dips its head into a hole) a hole not connected to the escape cage or the target hole without descending into the escape cage. Probe test for the cued learning was conducted one day after the last acquisition trial whereas mice trained in the spatial version of the test were submitted to 2 probe tests held 2 and 7 days after training completion (on days 8 and 13, respectively). In order to prevent memory extinction, animals were given three learning trials after the first probe test (data not shown). The target hole was disconnected from the escape cage and the mouse was given 5 min to explore the maze during which numbers of visits to each hole were counted.

*Object location*: This task evaluates spatial hippocampal-dependent memory and relies on the ability of rodents to discriminate between a novel and a familiar spatial location. Performances in this task are sensitive to the depletion of hippocampal adult neurogenesis.[11] This test takes place in a circular arena (40 cm diameter) containing a visual cue (rectangular striped pattern), surrounded by a white curtain, and is conducted in three phases: familiarization, exploration and test. One day before acquisition, each mouse was familiarized to the empty arena during 10 min. The next day, two identical objects are placed in the middle of the arena 20 cm apart. The mice are allowed to explore for 10 min during which the time spent sniffing the two objects was recorded. Mice that did not reach the criterion of 25 seconds of cumulated exploration for both objects were excluded. As a result, one mouse of each genotype was excluded from the analysis. On the test phase held one day later, copies of the same objects were exposed and one of the two objects was moved to a novel location. The position (left or right) of the displaced object was randomized to reduce bias towards a particular location. Mice were allowed to explore the objects during 10 min and the preference index for the displaced object was calculated as described below.

*Object recognition*: This task evaluates hippocampal-independent memory and assesses the ability of rodents to discriminate between a familiar and a novel object.[12, 13] Performances in this task are not sensitive to the depletion of hippocampal adult neurogenesis.[11, 12] A setup similar to the one described for the novel object location test was used, but the pattern was removed. The familiarization phase was identical to the object location task. The next day, mice were allowed to explore two identical objects placed in the center of the arena. The object exploration criterion was the same than for the object location task. Recognition memory was tested the following day. Mice were reintroduced for 10 min in the arena containing one familiar object and one novel object (left or right counterbalanced) which positions were identical to the acquisition phase. The preference for the novel object was calculated as described below.

*Preference index*: Object exploration was defined as the time spent actively sniffing or interacting with the object within a distance of 2 cm maximum. Measurement of the time spent exploring the displaced or novel object was expressed as a percentage of time spent exploring the displaced or novel object, related to the total exploration time for both objects during the test phase (in %, preference index; 50% is chance level).

*Metric pattern separation test*: This task evaluates memory processes that strongly rely on the dentate gyrus. Our protocol is based on the one described by Hunsaker[14] et al (2009). The setup consisted of a rectangular arena (60 x 40 x 30 cm) devoid of visual patterns. On the first day, mice were allowed to explore the empty arena for 20 minutes in groups of 4 mice, followed by 5 minutes of individual exploration. The next day, each mouse was allowed to explore for 15 min two different objects, placed 40 cm apart in the arena (exposition phase) during which exploration was measured by blocks of 5 min. Then, the mouse was placed in a waiting cage (standard cage with bedding material) during 20 min, and the objects were repositioned at a 10 cm distance. The mouse was then put back into the arena for a 5 min (test phase), during which it was free to re-explore the objects in their new metric configuration. Mice that did not reach the criterion of 20 seconds of total exploration time for both objects during the sample phase were removed from the experiment. Mice performances were evaluated *via* an exploration ratio calculated as follows: (exploration during 5 min of test phase) / (exploration during 5 min of test phase + exploration during the last 5 min of exposition phase). This constrained all the values between

0 and 1. Thus, an increased exploration during test phase was reflected by a ratio > 0.5, while a decreased exploration (or habituation) was reflected by a ratio < 0.5. Increased exploration of the new configuration demonstrates the ability of the mice to detect metric changes in the relationships between objects, as described by Hunsaker[14] et al, 2009.

#### Statistics

Data analysis was performed with Prism software (GraphPad.9). Morphological data analyses were evaluated with unpaired t-test, or Mann-Whitney test if the distribution data was not normal to compare parameters between groups. Behavioral analyses were evaluated with one sample t-test to compare preference index with chance level 50% or with 0.5 ratio and unpaired t-test to compare between genotypes. Repeated measurement two-way ANOVA with Bonferroni *post hoc* analyses were used when allowed. Data are presented as the mean ± SEM. Threshold for significance was set at $p<0.05$.

#### Data availability

The data that support the findings of this study are available from the corresponding author, upon reasonable request.
